# Phylogenetic and structural diversity of aromatically dense pili from environmental metagenomes

**DOI:** 10.1101/668343

**Authors:** M. S. Bray, J. Wu, C.C. Padilla, F. J. Stewart, D. A. Fowle, C. Henny, R. L. Simister, K. J. Thompson, S. A. Crowe, J. B. Glass

## Abstract

Electroactive type IV pili, or e-pili, are used by some microbial species for extracellular electron transfer. Recent studies suggest that e-pili may be more phylogenetically and structurally diverse than previously assumed. Here, we used updated aromatic density thresholds (≥9.8% aromatic amino acids, ≤22-aa aromatic gaps, and aromatic amino acids at residues 1, 24, 27, 50 and/or 51, and 32 and/or 57) to search for putative e-pilin genes in metagenomes from diverse ecosystems with active microbial metal cycling. Environmental putative e-pilins were diverse in length and phylogeny, and included truncated e-pilins in *Geobacter* spp., as well as longer putative e-pilins in Fe(II)-oxidizing *Betaproteobacteria* and *Zetaproteobacteria*.

**Originality and Significance:** Electroactive pili (e-pili) are used by microorganisms to respire solid metals in their environment through extracellular electron transfer. Thus, e-pili enable microbes to occupy specific environmental niches. Additionally, e-pili have important potential for biotechnological applications. Currently the repertoire of known e-pili is small, and their environmental distribution is largely unknown. Using sequence analysis, we identified numerous genes encoding putative e-pili from diverse anoxic, metal-rich ecosystems. Our results expand the diversity of putative e-pili in environments where metal oxides may be important electron acceptors for microbial respiration.

## Introduction

Electroactive microbes transport electrons through cell membranes into the extracellular environment (Sydow et al., 2014; Koch and Harnisch, 2016; Logan et al., 2019). These microbes play important roles in biogeochemical cycles in soils and sediments, bioremediation of toxic metals, and energy generation in microbial fuel cells (Lovley, 1991; Lovley and Coates, 1997; Logan, 2009; Lovley, 2011; Mahadevan et al., 2011). Electroactive *Deltaproteobacteria* in the genus *Geobacter* (order *Desulfuromonadales*) perform long-range extracellular electron transfer (EET) through electroactive pili (e-pili), composed of e-pilin structural subunits (Lovley, 2017; Lovley and Walker, 2019). *Geobacter* use e-pili for Fe(III) respiration, direct interspecies electron transfer (DIET), and growth on anodes (Reguera et al., 2005; Reguera et al., 2006; Rotaru et al., 2014).

*Geobacter* e-pili belong to the larger family of type IV-a pilins (T4aPs), which are broadly distributed in Bacteria and Archaea (Imam et al., 2011; Giltner et al., 2012; Berry and Pelicic, 2015). T4aPs have evolved to perform diverse cellular functions, including twitching motility, attachment, and genetic transformation. Most characterized *Geobacter* e-pilins are truncated versions of canonical T4aPs (Holmes et al., 2016). Type II (or “pseudopilin”) proteins are structurally similar to, but phylogenetically distinct from T4aPs, and assemble into type II secretion (T2S) systems instead of pili (Ayers et al., 2010).

Aromatic amino acid density seems to be essential for efficient electron transport in e-pili (Vargas et al., 2013; Liu et al., 2014; Liu et al., 2019). The close packing of aromatic residues within the pilus likely facilitates EET (Reardon and Mueller, 2013; Feliciano et al., 2015; Lovley, 2017). In particular, Phe1, Tyr24, and Tyr27 are key residues (Xiao et al., 2016), and Tyr32, Phe51 and Tyr57 also play important roles (Liu et al., 2019). The most conductive e-pilus measured to date is that of *Geobacter metallireducens*, which contains pilins that are 59 aa in mature length (after signal peptide sequence removal at the prepilin cleavage site) and comprised of 15.3% aromatics and no aromatic-free gaps >22 aa **(Table S1)**. The *G. metallireducens* e-pilus is 5000 times more conductive than the *Geobacter sulfurreducens* e-pilus, which has pilins which are 61 aa in mature length and comprised of 9.8% aromatics and no aromatic-free gaps >22 aa (Tan et al., 2017). The *G. sulfurreducens* e-pilus is 100 times more conductive than the *Geobacter uraniireducens* pilus, which contains much longer pilins (193 aa), 9.1% aromatics, and a 53 aa aromatic-free gap (Tan et al., 2016). Non-electroactive T4aPs are thought to be incapable of electroactivity due to insufficient aromatic residue packing (Feliciano et al., 2015; Malvankar et al., 2015; Kolappan et al., 2016). To our knowledge, the most aromatic-rich predicted e-pilus belongs to *Desulfobacula phenolica* (16.9%; Holmes et al., 2016).

Multiheme cytochromes (MHCs) are also involved in EET. Outer membrane MHCs move electrons from the periplasm into the extracellular environment (Aklujkar et al., 2013). The hexaheme OmcS can localize with *Geobacter* e-pili (Leang et al., 2010; Vargas et al., 2013; Liu et al., 2014). Conductive filaments comprised solely of OmcS were recovered from outer-membrane preparations of *G. sulfurreducens* grown in microbial fuel cells (Filman et al., 2019; Wang et al., 2019), but substantial evidence suggests that e-pilins in wild-type *Geobacter* cultures are comprised of PilA (Lovley and Walker, 2019).

Recently, the phylogenetic and structural diversity of e-pili has expanded beyond *Geobacter* spp. with the discovery of strongly conductive pili in clades outside of *Geobacter* genera, including *Syntrophus aciditrophicus* (*Deltaproteobacteria/Syntrophobacterales*), *Desulfurivibrio alkaliphilus* (*Deltaproteobacteria/Desulfobacterales*), *Calditerrivibrio nitroreducens* (*Deferribacteres*), and the archaeon *Methanospirillum hungatei* (*Euryarchaeota/Methanomicrobiales*) (Walker et al., 2018; Walker et al., 2019a; Walker et al., 2019b) **(Table 1)**. Pilin genes in these four microbes are much longer (110-182 aa) than in *Geobacter* spp., but have similar aromaticity (11-13%) and similar maximum aromatic-free gaps (22-35 aa). Pili from *Desulfofervidus auxilii*, *Shewanella oneidensis*, and *Pseudomonas aeruginosa* with minimal conductance have lower aromaticity (5.6-6.8%) and larger aromatic-free gaps (42-52 aa; Reguera et al., 2005; Liu et al., 2014; Walker et al., 2018). Therefore, it seems that aromatic density, defined here as percentage of aromatic amino acids and spacing of aromatic residues in the pilin sequence, is the key factor for identifying putative e-pilins based on sequence similarity (Walker et al., 2019a). In this study, we searched metagenomes from metal-rich environments and enrichment cultures for putative e-pilins based on aromatic density and spacing.

## Results

### Aromatic density and spacing distinguishes e-pilins from non-conductive T4aPs

We obtained published sequences for seven biochemically confirmed e-pilins, four non-conductive pilins (**Table S1**), and 35 functionally verified attachment/motility/competence T4aPs (**Table S2**). Biochemically confirmed e-pilins had mature lengths of 59-182 aa, 9.8-16.9% aromatics, and maximum aromatic-free gaps of 22-35 aa **(Figure 1; Table S1)**. Pilins implicated in functions other than long-range EET had 93-208 aa mature lengths, 3.5-11.0% aromatics, and 22-75 aa aromatic-free gaps **(Figure 1; Table S2)**. Sequence alignments showed that all bacterial e-pilins contained Phe1, Tyr24, Tyr27, and Tyr/Phe51. Most also contained an aromatic amino acid (Tyr or Phe) at residues 32, 50, and 57. Therefore, we used ≥9.8% aromatics, ≤22-aa aromatic-free gap, and the presence of aromatic amino acids at residues 1, 24, 27, 50 and/or 51, and 32 and/or 57 as a conservative threshold for predicting putative e-pilins from metagenomes, consistent with thresholds established by Walker et al. (2019a). Using these thresholds, two T4aPs in **Table S2** were predicted to be conductive: *G. sulfurreducens* OxpG, which forms a T2S system required for reduction of insoluble Fe(III) (Mehta et al., 2006), and *Dichelobacter nodosus* PilE, which is required for extracellular protease secretion and competence (Han et al., 2007).

**Figure 1.**
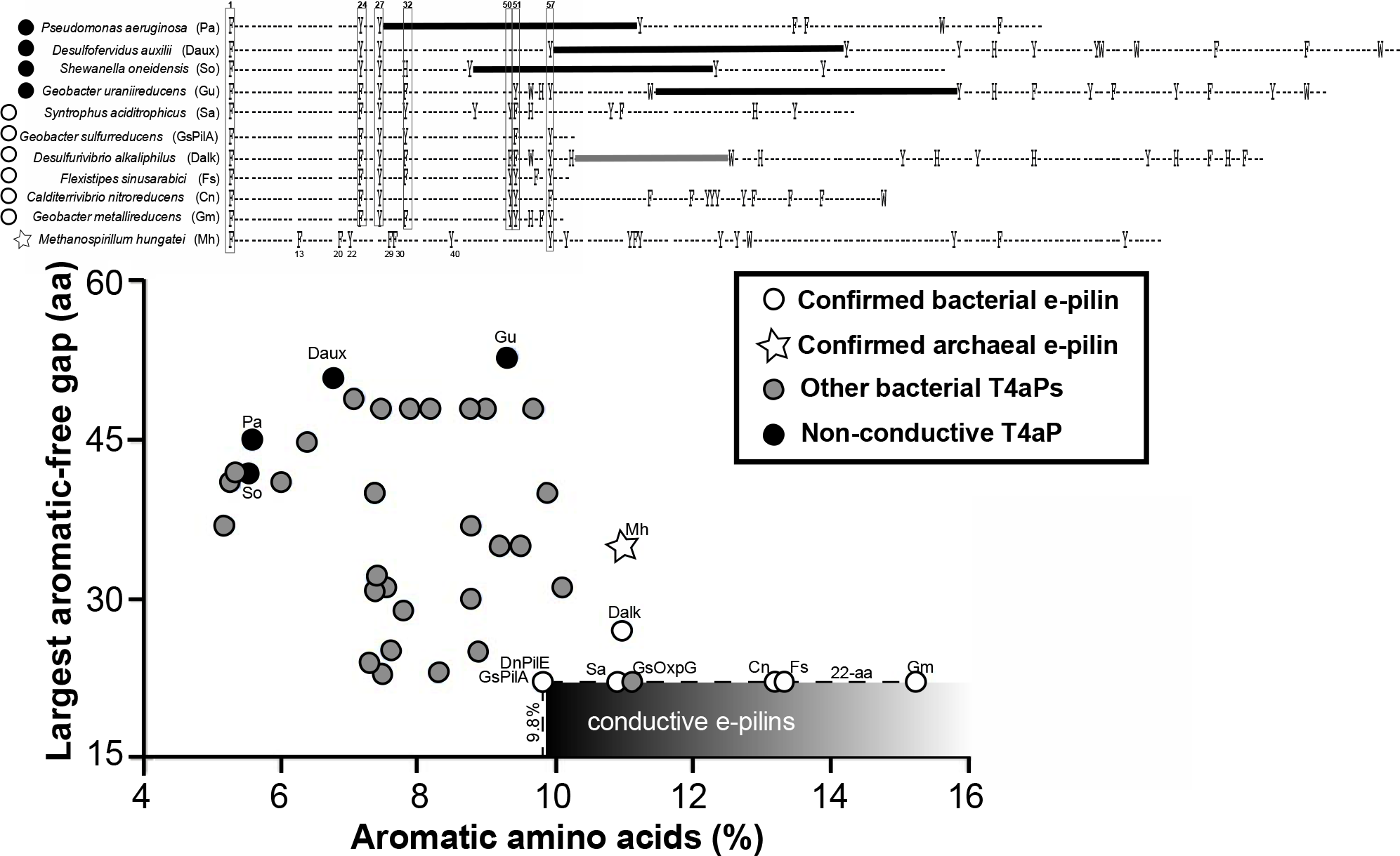
Basis for distinguishing e-pilins from other pilins. *Top:* Alignment showing the location of aromatic residues in each pilin tested for conductivity in previous studies (**Table S1**). Dark horizontal lines indicate 42-53 aa aromatic-free gaps in non-conductive pilins. Conserved N-terminal aromatic residues in bacterial e-pilins are indicated by vertical boxes. All bacterial e-pilins contained F-1, Y-24, Y-27, Y/F-51, Y/F-50 and/or Y/F-51, and H/Y/F-32 and/or Y/F-57. The only N-terminal residues shared by archaeal and bacterial e-pilins were F-1 and Y-57. *Bottom:* Relationship between gap size and percentage of aromatic amino acids in the mature pilin peptide for four types of pilins, which was used to establish conservative criteria for identifying putative e-pilins in environmental metagenomes. Therefore, we used ≥9.8% aromatics and ≤22-aa aromatic-free gap (boxed area labeled “conductive e-pilins”), and the presence of aromatic amino acids at residues as a conservative threshold for predicting putative e-pilins from metagenomes, consistent with thresholds established by Walker et al. (2019a). Additional information about each pilin is in **Tables S1** and **S2**.

### Putative e-pilins are present in ferruginous environments

We used the *G. sulfurreducens* e-pilin to query metagenomic contigs or metagenome-assembled genomes (MAGs) from environments with conditions amenable to metal respiration. We included metagenomes from ferruginous sediments from two lakes, Lake Matano and Lake Towuti, in the Malili Lakes system on Sulawesi, Indonesia, and the ferruginous water column from Kabuno Bay, Lake Kivu, Democratic Republic of Congo. These permanently stratified tropical lakes host one of the largest ferruginous environments on modern Earth with abundant iron-cycling microbes likely capable of EET (Crowe et al., 2007; Vuillemin et al., 2016). Other environments included deep groundwaters from Sweden (Asop Hard Rock), Japan (Horonobe Underground Laboratory), USA (Rifle, Colorado), and the North Atlantic (North Pond marine aquifer). We also included putative e-pilins from year-long laboratory incubations inoculated with Lake Matano sediment amended with Fe(III) or Mn(III) (see **Experimental Procedures**).

We screened the retrieved amino acid sequences for T4Ps using Pilfind (Imam et al., 2011), and the aromatic density thresholds established above (≥9.8% aromatic amino acids, ≤22-aa aromatic gaps, and aromatic amino acids at residues 1, 24, 27, 50 and/or 51, and 32 and/or 57). After partial sequences were removed, we recovered putative e-pilins ranging from 58 to 162 aa mature length with 9.8-15.5% aromatic density (**Table S3; Supplemental Data File**).

### Widening the phylogenetic diversity of putative e-pilins

To determine the phylogenetic diversity of environmental e-pilins, we constructed a maximum likelihood tree from an alignment of the T4aP amino acid sequences described above, as well as additional predicted *Deltaproteobacteria* e-pilins from cultured species (Holmes et al, 2016; Walker et al., 2018a) and BLAST searches **(Figure 2)**. *M. hungatei* e-pilin was used as the outgroup. The T4aP phylogeny was broadly consistent with previous findings (Holmes et al., 2016; Walker et al., 2018). All truncated e-pilins and all confirmed bacterial e-pilins clustered with *Deltaproteobacteria*. Non-conductive *Gammaprotebacteria* pilins and T2S pseudopilins fell on separate branches. Truncated *Desulfuromonadales* e-pilins (~60 aa) formed their own branch within the *Deltaproteobacteria* cluster. Other branches on the *Deltaproteobacteria* cluster contained recently discovered e-pilins from *Desulfobacterales*, *Deferribacteres*, and *Syntrophobacterales*. Roughly half of environmental putative e-pilins clustered with *Deltaproteobacteria*, including two putative e-pilins from native Lake Matano sediment and six putative e-pilins from >1 year anoxic incubations of Lake Matano sediments with Fe(III) oxides (**Table S3; Supplemental Data File).** Putative e-pilins from marine *Zetaproteobacteria* (*Mariprofundus micogutta* and two MAGs from the North Pond marine subsurface aquifer) and *Nitrospinae* (Crystal Geyser, Utah, USA) also clustered with *Deltaproteobacteria* e-pilins.

**Figure 2.**
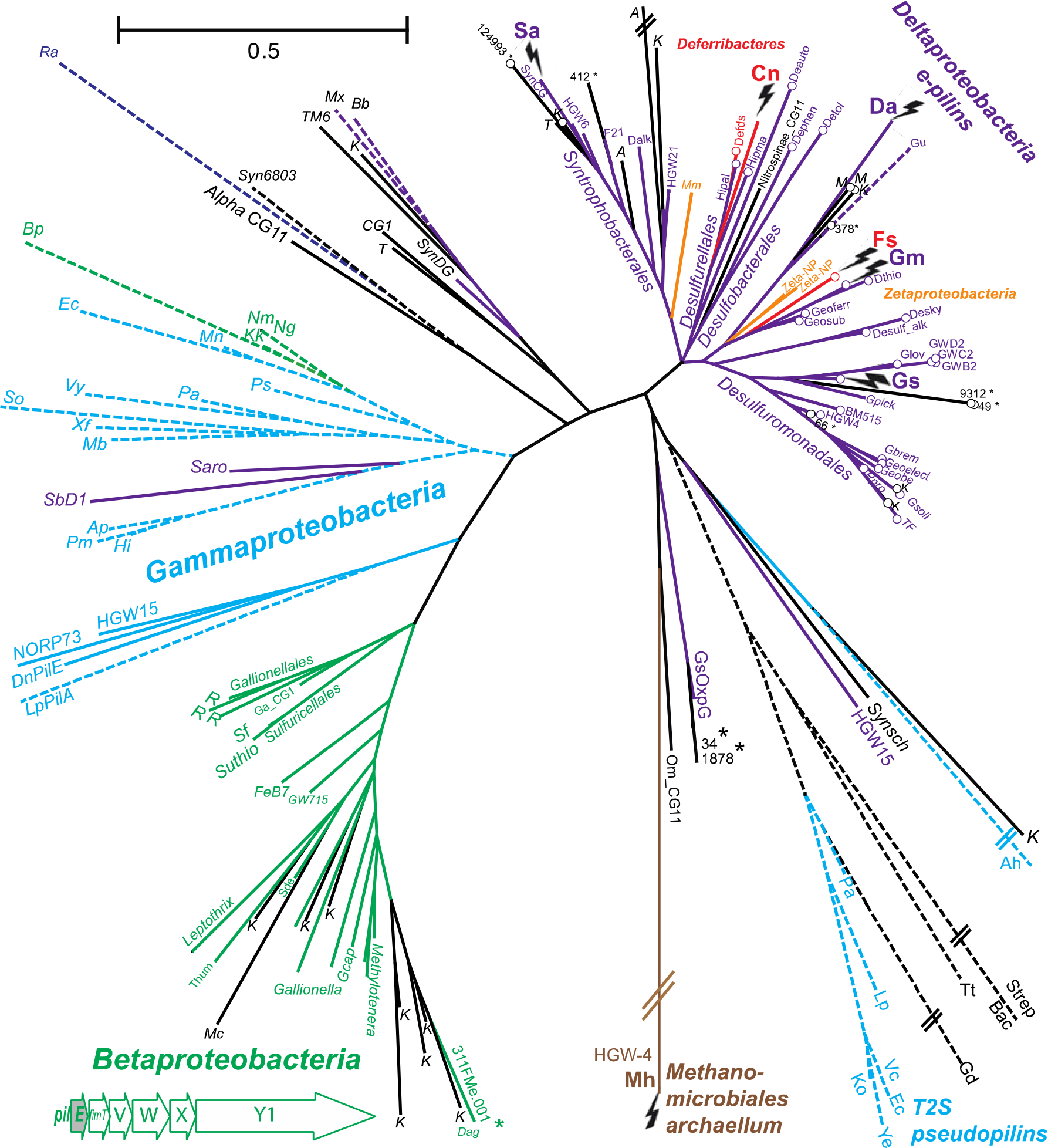
Maximum likelihood phylogenetic tree of pilin sequences. *Methanomicrobiales* archaellum was used as the outgroup. Asterisks indicate putative e-pilins from enrichment cultures. Dashed lines indicate sequences that do not meet the criteria for e-pili. Double lines indicate that the branch length was shortened to fit inside the figure boundaries. The gene order for *Betaproteobacteria* putative e-pilins (*pilE*) is shown. See **Supplemental Data File** for details on each sequence and full names of species. **Table S2** provides characteristics of type IV pilins and Type II secretion (T2S) pseudopilins involved in functions other than EET. Environmental abbreviations: A:Asop, CG:Crystal Geyser, K:Kabuno, M:Matano, Mc:McNutt, R:Rifle, T:Towuti.

Approximately half of environmental putative e-pilins fell outside the *Deltaproteobacteria* cluster on the T4aP phylogeny **(Figure 2)**. Eight unique putative e-pilin sequences (found 29 times in Kabuno Bay metagenomes), and one e-pilin from McNutt Creek (Georgia, USA), formed a distinct phylogenetic cluster with *pilE* genes from cultured *Betaproteobacteria* (*Gallionella, Leptothrix, Methylotenera, Sulfuricella, Thauera,* and *Dechloromonas*)*, Gallionellales* MAGs from groundwater, and *Rhodocyclales* MAG from Lake Matano enrichment cultures (311FMe.001; NCBI genome accession VAUH01000000). *Betaproteobacteria* PilE sequences in this clade contained 10.1-13.5% aromatics, ≤22-aa aromatic-free gaps, and key aromatic residues at positions 1, 24, 27, 50, 51, and 57. In all cases, putative *Betaproteobacteria pilE* genes were followed by *fimT-pilVWXY1*, which encode minor pilin assembly proteins (Nguyen et al., 2015).

Several putative environmental e-pilins clustered with non-conductive pilins from *Deltaproteobacteria, Gammaproteobacteria*, and *Firmicutes*. These included putative e-pilins in MAGs belonging to the candidate phylum *Dependentiae* (formerly TM6) from Rifle groundwater, *Alteromonas* NORP73 from North Pond marine subsurface aquifer, *Gammaproteobacteria* HGW15 from Horonobe Underground Laboratory, and *Proteobacteria* CG-11 from Crystal Geyser. The *G. sulfurreducens* OxpG, two sequences from Lake Matano enrichment cultures, and *Omnitrophica* sequences from Crystal Geyser were located on the same branch as the outgroup. *Omnitrophica* have been implicated in anaerobic respiration with metals (Hernsdorf et al., 2017) or sulfite (Anantharaman et al., 2018). To assess potential capacity for metal reduction, we searched MAGs that contained putative e-pilins for outer membrane/extracellular MHCs. Notably, the *Omnitrophica* MAG contained ten putative MHCs located adjacent to each other in the genome, three of which were predicted to be extracellular or outer-membrane MHCs, each with 11 or 13 hemes (**Figure S1**).

## Discussion

We recovered genes that meet *in silico* requirements for conductivity based on aromatic density and spacing both inside and outside of the well-established *Deltaproteobacteria* cluster. Our phylogenetic analyses suggest that the *Deltaproteobacteria* e-pilin genes have undergone more extensive horizontal gene transfer (HGT) than previously known. Our results suggest that truncated e-pilins are limited to the *Deltaproteobacteria* cluster, whereas predicted e-pilins outside of *Deltaproteobacteria* were full-length. In addition to their previously recognized HGT to several *Deferribacteres* species (Holmes et al, 2016; Walker et al., 2018a), we found putative e-pilins that clustered with *Deltaproteobacteria* in MAGs from *Nitrospinae* and *Zetaproteobacteria. Nitrospinae* are chemoautotrophic nitrite oxidizers that have not, to our knowledge, previously been implicated in EET. *Zetaproteobacteria*, the dominant marine Fe(II) oxidizers, were known to possess *pilA* genes, but the gene products were previously classified as non-conductive because they are >100 aa in length (He et al., 2017). Given the recent discovery of conductive e-pili with >100 aa (Walker et al., 2018), the possible occurrence of e-pilins in *Zetaproteobacteria* such as *Mariprofundus micogutta* needs to be re-evaluated.

Outside of the *Deltaproteobacteria* cluster, several putative e-pilin genes clustered with non-conductive *Gammaproteobacteria* pilin. These included *Alteromonas* NORP73 from the North Pond marine subsurface aquifer and *Gammaproteobacteria* HGW15 from Horonobe Underground Laboratory. *Alteromonas* are known to reduce Fe(III) and form electroactive biofilms (Vandecandelaere et al., 2008), but have not previously, to our knowledge, been found to possess e-pilins. The findings suggest that non-conductive full-length pilins may be capable of evolving conductive properties, although this awaits experimental validation.

Putative e-pilins were also found associated with clades not previously known to possess e-pili. Kabuno Bay metagenomes contained abundant e-pilin sequences most similar to those found in metabolically diverse *Betaproteobacteria* genera, including *Gallionella, Leptothrix, Methylotenera, Sulfuricella, Thauera,* and *Dechloromonas*. These putative e-pilin genes were classified as *pilE* and were followed by genes involved in minor pilus assembly. Putative *Betaproteobacteria* e-pili genes were also found in other groundwater MAGs, including Crystal Geyser, where *Gallionellaceae* are among the most abundant bacteria (Probst et al., 2018).

While the aromatically dense pilins in this study met the bioinformatic thresholds for e-pili, it is possible that they are used for another function, such as DIET (Holmes et al., 2017; Walker et al., 2019a) or cellular detection of solid surfaces via electrical communication (Lovley, 2017). Evaluation of the conductivity of the putative e-pilins awaits testing by genetic complementation of Δ*pilA* in *G. sulfurreducens*, as in Walker et al. (2018).

## Conclusions

This study identified putative e-pilins in the environment using aromatic density and gaps as the predictive tool, building off of previous studies that established the conductivity of longer PilA proteins (Walker et al., 2018). The sequences we recovered suggest that e-pilins are both phylogenetically and structurally diverse. We conclude that e-pili may be composed of pilin monomers of a variety of lengths and aromatic densities, and that diverse bacteria, including Fe(II)-oxidizing *Betaproteobacteria* and *Zetaproteobacteria*, may use e-pili for EET or possibly other unknown functions.

## Acknowledgements

This research was funded by NASA Exobiology grant NNX14AJ87G with support from the NASA Astrobiology Institute (NNA15BB03A). SAC, RLS, and KTJ were supported through NSERC Discovery grant 0487, CFI projects 229652 and 36071, and the Canada Research Chairs program. We thank Bianca Costa, Miles Mobley, Benjamin Reed, and Johnny Striepen for assistance with laboratory incubations, and Sean Elliott and Betül Kaçar for helpful discussions.

## Experimental Procedures

### Sampling and enrichment of Lake Matano Sediment

Two sediment cores were obtained from 590 m water depth in Lake Matano, Sulawesi Island, Indonesia in May 2010 (2°28’S, 121°20’E, *in situ* sediment temperature ~27°C) and stored under anoxic conditions. The sediments were mixed with anoxic freshwater media in a 1:5 ratio in an anoxic chamber and dispensed in stoppered serum bottles, as in Bray et al. (2017). Cultures were amended first with goethite and later with ferrihydrite. They were incubated for 490 days at 30°C, with multiple transfers, each time diluting the original sediment with freshwater media. Sediment had been diluted over 1000-fold by the time DNA was extracted for sequencing. Details on metagenomes from 395-day anoxic enrichments of Lake Matano sediment incubated with Mn(III) pyrophosphate are reported in a separate publication (Szeinbaum et al., 2019).

### DNA extraction and metagenome sequencing, assembly, binning and annotation

Community DNA from Lake Matano sediment enrichments was extracted from 2◻g samples and purified using a PowerSoil Isolation Kit and UltraClean® 15 Purification Kit (formerly MO BIO Laboratories, now Qiagen, Carlsbad, CA, USA) following the manufacturer’s protocol. Indexed libraries were created from purified community DNA using the NexteraXT DNA Sample Prep kit (Illumina, San Diego, CA, USA) following manufacturer instructions. Libraries were pooled and sequenced on two runs of an Illumina MiSeq using a 500 cycle (paired end 250 × 250 bp) kit. Illumina reads were quality trimmed using Trim Galore! (http://www.bioinformatics.babraham.ac.uk/projects/trim_galore/) with a quality score and minimum length cutoff of Q25 and 100 bp, respectively, and merged with FLASH with the shortest overlap of 25 bp. Barcoded sequences were de-multiplexed, trimmed (length cutoff 100 bp), and filtered to remove low quality reads (average Phred score <25) using Trim Galore!. Forward and reverse reads were assembled using SPAdes (Nurk et al., 2013) with the ‘meta’ option. The number of contigs, contig length, GC content, N50, and L50 assembly statistics were calculated with metaQUAST (Mikheenko et al., 2015). Raw sequence reads, and all genomic bins were deposited in NCBI under the accession number <u>PRJNA505658</u>.

### E-pilin identification from microbial metagenomes

Environmental metagenomes and MAGs were downloaded from IMG-JGI and NCBI (see **Table 1** for taxon object IDs). For all metagenomes, Prodigal (Hyatt et al., 2010) was used to predict genes from contig files and write them to amino acid fasta files. Amino acid sequences from MAGs were downloaded directly from NCBI. Predicted protein files were then used as databases for protein BLAST, using the *G. sulfurreducens* PilA protein as query. Hits with a bit score greater than 55 were pulled from the databases. These recovered sequences were then further verified as T4P using Pilfind (http://signalfind.org/pilfind.html), a web tool that identifies type IV pilin signal sequences (Imam et al., 2011). Pilin amino acid sequences were then run through a python script that calculated the mature pilin length, percent aromatic amino acids, and aromatic free gaps (https://github.com/GlassLabGT/Python-scripts). Partial genes were retained if truncated on the N-terminus before the signal peptide and removed if truncated on the C-terminus. Remaining sequences were manually screened for the presence of aromatic amino acids at residues 1, 24, 27, 50 and/or 51, and 32 and/or 57.

### Pilin multiple sequence alignment and phylogenetic analysis

Identified pilin amino acid sequences were aligned using MUSCLE and a maximum likelihood tree was constructed using MEGA. The alignment is provided as **Supplemental Material**. The evolutionary history was inferred by using the Maximum Likelihood method based on the JTT matrix-based model (Jones et al., 1992). Archaeal pili from *M. hungatei* and two other *Methanomicrobiales* were used for the outgroup. The tree with the highest log likelihood is shown. Initial tree(s) for the heuristic search were obtained automatically by applying Neighbor-Join and BioNJ algorithms to a matrix of pairwise distances estimated using a JTT model, and then selecting the topology with superior log likelihood value. There were 52 positions total in the final dataset. Evolutionary analyses were conducted in MEGA7 (Kumar et al., 2016).

### Multiheme cytochrome analysis

Groundwater MAGs in which we identified aromatically dense pilins were further probed for the presence of multiheme cytochrome proteins. Amino acids files were run through the “cytochrome_stats.py” described in (Badalamenti et al., 2016) available at https://github.com/bondlab/scriptsm, which identifies proteins with 3 or more cytochrome-binding motifs (Cxx(x)CH).

## References

Aklujkar, M., Coppi, M., Leang, C., Kim, B., Chavan, M., Perpetua, L., Giloteaux, L., Liu, A., and Holmes, D. (2013) Proteins involved in electron transfer to Fe(III) and Mn(IV) oxides by *Geobacter sulfurreducens* and *Geobacter uraniireducens*. Microbiology 159: 515–535.

Anantharaman, K., Hausmann, B., Jungbluth, S.P., Kantor, R.S., Lavy, A., Warren, L.A., Rappé, M.S., Pester, M., Loy, A., and Thomas, B.C. (2018) Expanded diversity of microbial groups that shape the dissimilatory sulfur cycle. ISME J 12: 1715.

Ayers, M., Howell, P.L., and Burrows, L.L. (2010) Architecture of the type II secretion and type IV pilus machineries. Future Microbiol 5: 1203–1218.

Badalamenti, J.P., Summers, Z.M., Chan, C.H., Gralnick, J.A., and Bond, D.R. (2016) Isolation and genomic characterization of ‘*Desulfuromonas soudanensis* WTL’, a metal-and electrode-respiring bacterium from anoxic deep subsurface brine. Front Microbiol 7: 913.

Berry, J.-L., and Pelicic, V. (2015) Exceptionally widespread nanomachines composed of type IV pilins: the prokaryotic Swiss Army knives. FEMS Microbiol Rev 39: 134–154.

Bray, M.S., Wu, J., Reed, B.C., Kretz, C.B., Belli, K.M., Simister, R.L., Henny, C., Stewart, F.J., DiChristina, T.J., and Brandes, J.A. (2017) Shifting microbial communities sustain multi-year iron reduction and methanogenesis in ferruginous sediment incubations. Geobiology 15: 678–689.

Crowe, S.A., O’Neill, A.H., Kulczycki, E., Weisener, C.G., Roberts, J.A., and Fowle, D.A. (2007) Reductive dissolution of trace metals from sediments. Geomicrobiol J 24: 157–165.

Feliciano, G., Steidl, R., and Reguera, G. (2015) Structural and functional insights into the conductive pili of *Geobacter sulfurreducens* revealed in molecular dynamics simulations. Phys Chem Chem Phys 17: 22217–22226.

Filman, D.J., Marino, S.F., Ward, J.E., Yang, L., Mester, Z., Bullitt, E., Lovley, D.R., and Strauss, M. (2019) Cryo-EM reveals the structural basis of long-range electron transport in a cytochrome-based bacterial nanowire. Comm Biol 2: 219.

Giltner, C.L., Nguyen, Y., and Burrows, L.L. (2012) Type IV pilin proteins: Versatile molecular modules. Microbiol Mol Biol Rev 76: 740–772.

Han, X., Kennan, R.M., Parker, D., Davies, J.K., and Rood, J.I. (2007) Type IV fimbrial biogenesis is required for protease secretion and natural transformation in *Dichelobacter nodosus*. J Bacteriol 189: 5022–5033.

He, S., Barco, R.A., Emerson, D., and Roden, E.E. (2017) Comparative genomic analysis of neutrophilic iron (II) oxidizer genomes for candidate genes in extracellular electron transfer. Front Microbiol 8: 1584.

Hernsdorf, A.W., Amano, Y., Miyakawa, K., Ise, K., Suzuki, Y., Anantharaman, K., Probst, A., Burstein, D., Thomas, B.C., and Banfield, J.F. (2017) Potential for microbial H2 and metal transformations associated with novel bacteria and archaea in deep terrestrial subsurface sediments. ISME J 11: 1915–1929.

Holmes, D.E., Dang, Y., Walker, D.J., and Lovley, D.R. (2016) The electrically conductive pili of *Geobacter* species are a recently evolved feature for extracellular electron transfer. Microb Genom 2: e000072.

Holmes, D.E., Shrestha, P.M., Walker, D.J., Dang, Y., Nevin, K.P., Woodard, T.L., and Lovley, D.R. (2017) Metatranscriptomic evidence for direct interspecies electron transfer between Geobacter and Methanothrix species in methanogenic rice paddy soils. Appl Environ Microbiol 83: e00223–00217.

Hyatt, D., Chen, G.-L., LoCascio, P.F., Land, M.L., Larimer, F.W., and Hauser, L.J. (2010) Prodigal: prokaryotic gene recognition and translation initiation site identification. BMC Bioinformatics 11: 119.

Imam, S., Chen, Z., Roos, D.S., and Pohlschröder, M. (2011) Identification of surprisingly diverse type IV pili, across a broad range of gram-positive bacteria. PloS One 6: e28919.

Jones, D.T., Taylor, W.R., and Thornton, J.M. (1992) The rapid generation of mutation data matrices from protein sequences. Bioinformatics 8: 275–282.

Koch, C., and Harnisch, F. (2016) Is there a specific ecological niche for electroactive microorganisms? ChemElectroChem 3: 1282–1295.

Kolappan, S., Coureuil, M., Yu, X., Nassif, X., Egelman, E.H., and Craig, L. (2016) Structure of the *Neisseria meningitidis* type IV pilus. Nature Comm 7: 13015.

Kumar, S., Stecher, G., and Tamura, K. (2016) MEGA7: Molecular Evolutionary Genetics Analysis version 7.0 for bigger datasets. Mol Biol Evol 33: 1870–1874.

Leang, C., Qian, X., Mester, T., and Lovley, D.R. (2010) Alignment of the c-type cytochrome OmcS along pili of *Geobacter sulfurreducens*. Appl Environ Microbiol 76: 4080–4084.

Liu, X., Tremblay, P.-L., Malvankar, N.S., Nevin, K.P., Lovley, D.R., and Vargas, M. (2014) A *Geobacter sulfurreducens* strain expressing *Pseudomonas aeruginosa* type IV pili localizes OmcS on pili but is deficient in Fe (III) oxide reduction and current production. Appl Environ Microbiol 80: 1219–1224.

Liu, X., Wang, S., Xu, A., Zhang, L., Liu, H., and Ma, L.Z. (2019) Biological synthesis of high-conductive pili in aerobic bacterium *Pseudomonas aeruginosa*. Appl Micro Biotechnol 103: 1535–1544.

Logan, B.E. (2009) Exoelectrogenic bacteria that power microbial fuel cells. Nat Rev Microbiol 7: 375–381.

Logan, B.E., Rossi, R., and Saikaly, P.E. (2019) Electroactive microorganisms in bioelectrochemical systems. Nat Rev Microbiol 17: 307–319.

Lovley, D.R. (1991) Dissimilatory Fe(III) and Mn(IV) reduction. Microbiol Rev 55: 259–287.

Lovley, D.R. (2011) Live wires: direct extracellular electron exchange for bioenergy and the bioremediation of energy-related contamination. Energ Environ Sci 4: 4896–4906.

Lovley, D.R. (2017) Electrically conductive pili: biological function and potential applications in electronics. Curr Opin Electrochem 4: 190–198.

Lovley, D.R., and Coates, J.D. (1997) Bioremediation of metal contamination. Curr Opin Biotechnol 8: 285–289.

Lovley, D.R., and Walker, D.J.F. (2019) *Geobacter* protein nanowires. PeerJ Preprints 7: e27773v27771.

Mahadevan, R., Palsson, B.Ø., and Lovley, D.R. (2011) *In situ* to *in silico* and back: elucidating the physiology and ecology of *Geobacter* spp. using genome-scale modelling. Nat Rev Microbiol 9: 39–50.

Malvankar, N.S., Vargas, M., Nevin, K., Tremblay, P.-L., Evans-Lutterodt, K., Nykypanchuk, D., Martz, E., Tuominen, M.T., and Lovley, D.R. (2015) Structural basis for metallic-like conductivity in microbial nanowires. mBio 6: e00084–00015.

Mehta, T., Childers, S.E., Glaven, R., Lovley, D.R., and Mester, T. (2006) A putative multicopper protein secreted by an atypical type II secretion system involved in the reduction of insoluble electron acceptors in *Geobacter sulfurreducens*. Microbiology 152: 2257–2264.

Mikheenko, A., Saveliev, V., and Gurevich, A. (2015) MetaQUAST: evaluation of metagenome assemblies. Bioinformatics 32: 1088–1090.

Nguyen, Y., Sugiman-Marangos, S., Harvey, H., Bell, S.D., Charlton, C.L., Junop, M.S., and Burrows, L. L. (2015) *Pseudomonas aeruginosa* minor pilins prime type IVa pilus assembly and promote surface display of the PilY1 adhesin. J Biol Chem 290: 601–611.

Nurk, S., Bankevich, A., Antipov, D., Gurevich, A.A., Korobeynikov, A., Lapidus, A., Prjibelski, A.D., Pyshkin, A., Sirotkin, A., and Sirotkin, Y. (2013) Assembling single-cell genomes and mini-metagenomes from chimeric MDA products. J Comp Biol 20: 714–737.

Probst, A.J., Ladd, B., Jarett, J.K., Geller-McGrath, D.E., Sieber, C.M., Emerson, J.B., Anantharaman, K., Thomas, B.C., Malmstrom, R.R., and Stieglmeier, M. (2018) Differential depth distribution of microbial function and putative symbionts through sediment-hosted aquifers in the deep terrestrial subsurface. Nature Microbiol 3: 328–336.

Reardon, P.N., and Mueller, K.T. (2013) Structure of the type IVa major pilin from the electrically conductive bacterial nanowires of *Geobacter sulfurreducens*. J Biol Chem 288: 29260–29266.

Reguera, G., McCarthy, K.D., Mehta, T., Nicoll, J.S., Tuominen, M.T., and Lovley, D.R. (2005) Extracellular electron transfer via microbial nanowires. Nature 435: 1098–1101.

Reguera, G., Nevin, K.P., Nicoll, J.S., Covalla, S.F., Woodard, T.L., and Lovley, D.R. (2006) Biofilm and nanowire production leads to increased current in *Geobacter sulfurreducens* fuel cells. Appl Environ Microbiol 72: 7345–7348.

Rotaru, A.-E., Shrestha, P.M., Liu, F., Markovaite, B., Chen, S., Nevin, K.P., and Lovley, D.R. (2014) Direct interspecies electron transfer between *Geobacter metallireducens* and *Methanosarcina barkeri*. Appl Environ Microbiol 80: 4599–4605.

Sydow, A., Krieg, T., Mayer, F., Schrader, J., and Holtmann, D. (2014) Electroactive bacteria—molecular mechanisms and genetic tools. Appl Micro Biotechnol 98: 8481–8495.

Szeinbaum, N., Nunn, B.L., Cavazos, A.R., Crowe, S.A., Stewart, F.J., DiChristina, T.J., Reinhard, C.T., and Glass, J.B. (2019) Expression of extracellular multiheme cytochromes discovered in a betaproteobacterium during Mn(III) reduction. BioRXiv: 695007.

Tan, Y., Adhikari, R.Y., Malvankar, N.S., Ward, J.E., Woodard, T.L., Nevin, K.P., and Lovley, D.R. (2017) Expressing the *Geobacter metallireducens* PilA in *Geobacter sulfurreducens* yields pili with exceptional conductivity. mBio 8: e02203–02216.

Tan, Y., Adhikari, R.Y., Malvankar, N.S., Ward, J.E., Nevin, K.P., Woodard, T.L., Smith, J.A., Snoeyenbos-West, O.L., Franks, A.E., and Tuominen, M.T. (2016) The low conductivity of *Geobacter uraniireducens* pili suggests a diversity of extracellular electron transfer mechanisms in the genus *Geobacter*. Front Microbiol 7: 980.

Vandecandelaere, I., Nercessian, O., Segaert, E., Achouak, W., Mollica, A., Faimali, M., De Vos, P., and Vandamme, P. (2008) *Alteromonas genovensis* sp. nov., isolated from a marine electroactive biofilm and emended description of *Alteromonas macleodii* Baumann et al. 1972 (Approved Lists 1980). Int J Syst Evol Microbiol 58: 2589–2596.

Vargas, M., Malvankar, N.S., Tremblay, P.-L., Leang, C., Smith, J.A., Patel, P., Synoeyenbos-West, O., Nevin, K.P., and Lovley, D.R. (2013) Aromatic amino acids required for pili conductivity and long-range extracellular electron transport in *Geobacter sulfurreducens*. mBio 4: e00105–00113.

Vuillemin, A., Friese, A., Alawi, M., Henny, C., Nomosatryo, S., Wagner, D., Crowe, S.A., and Kallmeyer, J. (2016) Geomicrobiological features of ferruginous sediments from Lake Towuti, Indonesia. Front Microbiol 7: 1007.

Walker, D.J., Nevin, K.P., Holmes, D.E., Rotaru, A.-E., Ward, J.E., Woodard, T.L., Zhu, J., Ueki, T., Nonnenmann, S.S., and McInerney, M.J. (2019a) *Syntrophus* conductive pili demonstrate that common hydrogen-donating syntrophs can have a direct electron transfer option. BioRXiv: 479683.

Walker, D.J.F., Martz, E., Holmes, D.E., Zhou, Z., Nonnenmann, S.S., and Lovley, D.R. (2019b) The archaellum of *Methanospirillum hungatei* is electrically conductive. mBio 10: e00579–00519.

Walker, D.J.F., Adhikari, R.Y., Holmes, D.E., Ward, J.E., Woodard, T.L., Nevin, K.P., and Lovley, D.R. (2018) Electrically conductive pili from pilin genes of phylogenetically diverse microorganisms. ISME J 12: 48–58.

Wang, F., Gu, Y., O’Brien, J.P., Sophia, M.Y., Yalcin, S.E., Srikanth, V., Shen, C., Vu, D., Ing, N.L., and Hochbaum, A.I. (2019) Structure of microbial nanowires reveals stacked hemes that transport electrons over micrometers. Cell 177: 361–369. e310.

Xiao, K., Malvankar, N.S., Shu, C., Martz, E., Lovley, D.R., and Sun, X. (2016) Low energy atomic models suggesting a pilus structure that could account for electrical conductivity of *Geobacter sulfurreducens* pili. Sci Rep 6: 23385.

