## Supplemental Material for "Phylogenetic and structural diversity of aromatically dense pili from environmental metagenomes"

Running Title: Predicting e-pili

^#^Now at: Dovetail Genomics, LLC, Santa Cruz, CA, US

| **GenBank Accession**  **or Locus Tag** | **Species and Abbreviation** | **Mature**  **length (aa)** | **% aromatics** | **Maximum aromatic-free gap (aa)** | **Conductive? Y: yes; N: no;**  **U: unmeasured** | **Reference** |
| --- | --- | --- | --- | --- | --- | --- |
| P04739 | *Pseudomonas aeruginosa* (Pa) | 143 | 5.6 | 45 | N | Liu et al. (2019) |
| WP_011070777 | *Shewanella oneidensis* (So) | 126 | 5.6 | 42 | N | Reguera et al. (2005) |
| AMM39924 | *Desulfofervidus auxilii* (Daux) | 206 | 6.8 | 51 | N | (Walker et al., 2018) |
| Gura_2677 | *Geobacter uraniireducens* (Gu) | 193 | 9.1 | 53 | N | Tan et al. (2016) |
| **GSU1496** | ***Geobacter sulfurreducens* (GsPilA)** | **61** | **9.8** | **22** | **Y** | **Reguera et al. (2005)** |
| Glov_2096 | *Geobacter lovleyi* (Glov) | 60 | 10.0 | 22 | U |  |
| KN63_04755 | *Smithella* sp. F21 (F21) | 109 | 10.1 | 22 | U |  |
| WP_028893309 | *Syntrophorhabdus aromaticivorans* (Sar) | 128 | 10.2 | 27 | U |  |
| WP_084710848 | *Syntrophaceticus schinkii* (Synsch) | 105 | 10.5 | 25 | U |  |
| WP_039645155 | *Geobacter soli* (Gsoli) | 65 | 10.8 | 22 | U |  |
| Ppro_1656 | *Pelobacter propionicus* (Ppro) | 64 | 10.9 | 22 | U |  |
| WP_027714750 | *Desulfuromonas* sp. TF (TF) | 64 | 10.9 | 22 | U |  |
| WP_020676438 | *Geopsychrobacter electrodiphilus* (Geoelect) | 64 | 10.9 | 22 | U |  |
| **SYN_00814** | ***Syntrophus aciditrophicus* (Sa)** | **110** | **10.9** | **22** | **Y** | **Walker et al. (2019a)** |
| **DaAHT2_2283** | ***Desulfurivibrio alkaliphilus* (Da)** | **182** | **11.0** | **27** | **Y** | **Walker et al. (2018)** |
| **Mhun_3140** | ***Methanospirillum hungatei* (Mh)** | **164** | **11.0** | **35** | **Y** | **Walker et al. (2019b)** |
| SPF50166 | *Syntrophobacter* sp. SbD1 (SbD1) | 145 | 11.7 | 21 | U |  |
| WP_039742239 | *Geobacter pickeringii* (Gpick) | 60 | 11.7 | 22 | U |  |
| OR1_03834 | *Geobacter* sp. OR-1 | 64 | 12.5 | 22 | U |  |
| AMJ45_02600 | *Syntrophobacter* sp. DG_60 (SynDG) | 159 | 12.6 | 22 | U |  |
| WP_040097998 | *Geoalkalibacter ferrihydriticus* (Geoferr) | 61 | 13.1 | 22 | U |  |
| WP_022670383 | *Hippea alviniae* (Hipal) | 61 | 13.1 | 22 | U |  |
| WP_083610739 | *Desulfatibacillum alkenivorans* (Dalk) | 114 | 13.2 | 22 | U |  |
| WP_144681039 | *Desulfobotulus alkaliphilus* (Desulf_alk) | 60 | 13.3 | 22 | U |  |
| **Flexsi_2288** | ***Flexistipes sinusarabici* (Fs)** | **60** | **13.3** | **22** | **Y** | **Walker et al. (2018)** |
| **Calni_0149** | ***Calditerrivibrio nitroreducens* (Cn)** | **119** | **13.4** | **22** | **Y** | **Walker et al. (2018)** |
| Gbem_2590 | *Geobacter bemidjiensis* (Gbem) | 66 | 13.6 | 22 | U |  |
| Hipma_0737 | *Hippea maritima* (Hipma) | 59 | 13.6 | 22 | U |  |
| DEFDS_1270 | *Deferribacter desulfuricans* | 59 | 13.6 | 22 | U |  |
| WP_092080546 | *Desulfuromonas thiophila* (Dthio) | 59 | 13.6 | 22 | U |  |
| TOL2_21350 | *Desulfobacula toluolica* (Dtol) | 58 | 13.8 | 22 | U |  |
| WP_092349663 | *Desulfuromusa kysingii* (Desky) | 60 | 15.0 | 22 | U |  |
| HRM2_27700 | *Desulfobacterium autotrophicum* (Deauto) | 59 | 15.3 | 22 | U |  |
| **Gmet_1399** | ***Geobacter metallireducens*** (Gm) | **59** | **15.3** | **22** | **Y** | **Tan et al. (2017)** |
| WP_040199521 | *Geoalkalibacter subterraneus* (Geosub) | 66 | 15.5 | 22 | U |  |
| WP_026840453 | *Geobacter bremensis* (Gbrem) | 64 | 15.6 | 22s | U |  |
| WP_092237916 | *Desulfobacula phenolica* (Dephen) | 59 | 16.9 | 22 | U |  |

**Table S1.** **Characteristics of predicted or confirmed e-pilin proteins, and non-conductive pilins, in order of increasing aromatic density, from cultured isolates, based on Holmes et al. 2016 and Walker et al. 2018.** These proteins listed as “U” are predicted to infer long-range electron transport capability in the listed species, based on the criteria listed in the paper. Biochemically confirmed e-pilins are bolded.

| **GenBank**  **Accession** | **Species and Abbreviation** | **Protein name** | **Function** | **Mature**  **length** | **% aromatics** | **Maximum aromatic-free gap (aa)** | **Reference** |
| --- | --- | --- | --- | --- | --- | --- | --- |
| WP_004090380 | *Xylella fastidiosa* (Xf) | PilA | M | 142 | 3.5 | 75 | Li et al. (2007) |
| WP_011912242 | *Pseudomonas stutzeri* (Ps) | PilA | C | 162 | 4.9 | 66 | Graupner et al. (2000) |
| CAD14088 | *Ralstonia solanacearum* (Rs) | PilA | C, A, M | 154 | 5.2 | 37 | Kang et al. (2002) |
| P07640 | *Moraxella bovis* (Mb) | TfpQ | A | 151 | 5.3 | 41 | Marrs et al. (1985) |
| CAH34774 | *Burkholderia pseudomallei* (Bp) | PilA | A | 167 | 5.4 | 42 | Essex-Lopresti et al. (2005) |
| P04739 | *Pseudomonas aeruginosa* (PaPilA) | PilA | A, M | 143 | 5.6 | 45 | (Doig et al., 1988; Burrows, 2012; Liu et al., 2019) |
| WP_011070777 | *Shewanella oneidensis* (So) | PilA | A, M | 126 | 5.6 | 42 | Thormann et al. (2004) |
| WP_019390097 | *Kingella kingae* (Kk) | PilA | A | 149 | 6.0 | 41 | Kehl-Fie et al. (2008) |
| CAC94922 | *Ruminococcus albus* (Ra) | PilA | A | 157 | 6.4 | 45 | Rakotoarivonina et al. (2005) |
| WP_011035902 | *Xanthomonas campestris* (Xc) | XpsG | S | 126 | 7.1 | 49 | Nien-Tai et al. (2002) |
| WP_005694367 | *Haemophilus influenzae* (Hi) | PilA | C, M | 137 | 7.3 | 24 | (Dougherty and Smith, 1999; Bakaletz et al., 2005) |
| NP_273084 | *Neisseria meningitidis* (Nm) | PilE | A | 163 | 7.4 | 31 | Nassif et al. (1994) |
| AAA25310 | *Moraxella nonliquefaciens* (Mn) | PilA | A | 148 | 7.4 | 40 | Tønjum et al. (1991) |
| AAL32049 | *Vibrio vulnificus* (Vv) | PilA | A | 135 | 7.4 | 32 | Paranjpye and Strom (2005) |
| B657_24710 | *Bacillus subtilis* (Bac) | ComGC | C | 93 | 7.5 | 23 | Chen et al. (2006) |
| WP_010872381 | *Synechocystis* PCC 6803 (Syn6803) | HofG | M | 147 | 7.5 | 48 | Bhaya et al. (2000) |
| AAA25466 | *Neisseria gonorrhoeae* (Ng) | PilE | C, A | 159 | 7.5 | 31 | Rudel et al. (1992) |
| AAC28468 | *Eikenella corrodens* (Ec) | PilA | C, M | 145 | 7.6 | 25 | Villar et al. (2001) |
| WP_000738626 | *Streptococcus pneumoniae* (Strep) | ComGC | C | 102 | 7.8 | 29 | Laurenceau et al. (2013) |
| CAC83032 | *Yersinia enterocolitica* (Ye) | Yts1G | S | 139 | 7.9 | 48 | von Tils et al. (2012) |
| EUC91830 | *Klebsiella oxytoca* (Ko) | PulG | S | 134 | 8.2 | 48 | Pugsley (1993) |
| WP_005741096 | *Pasteurella multocida* (Pm) | PilA | A | 132 | 8.3 | 23 | Marrs et al. (1985) |
| AAM55486 | *Thermus thermophilus* (Tt) | PilA | C, M | 125 | 8.8 | 37 | Rumszauer et al. (2006) |
| WP_000738789 | *Vibrio cholerae* (Vc) | EspG | S | 137 | 8.8 | 30 | Sandkvist et al. (1997) |
| NP_417787 | *Escherichia coli* K12 (Ec) | GspG | S | 136 | 8.8 | 48 | Sauvonnet et al. (2000) |
| ABN73976 | *Actinobacillus pleuropneumoniae* (Ap) | ApfA | A | 135 | 8.9 | 25 | Zhang et al. (2000) |
| WP_011704371 | *Aeromonas hydrophila* (Ah) | ExeG | S | 202 | 8.9 | 25 | Howard et al. (1993) |
| WP_003103532 | *Pseudomonas aeruginosa* (Pa) | XcpT | S | 134 | 9.0 | 48 | Hahn (1997) |
| AAK58496 | *Gluconacetobacter diazotrophicus* (Gd) | LsdG | S | 130 | 9.2 | 35 | Arrieta et al. (2004) |
| CAE79186 | *Bdellovibrio bacteriovorus* (Bb) | PilA | A | 179 | 9.5 | 35 | Chanyi and Koval (2014) |
| AAD43220 | *Legionella pneumophila* (Lp) | LspG | S | 134 | 9.7 | 48 | Rossier et al. (2004) |
| WP_012031206 | *Dichelobacter nodosus** (DnPilE) | PilE | C, S, F | 143 | 9.8 | 22 | Han et al. (2007) |
| AAC38305 | *Legionella pneumophila* (LpPilA) | PilA | A | 132 | 9.9 | 40 | Stone and Kwaik (1998) |
| WP_011555734 | *Myxococcus xanthus* (Mx) | PilA | M | 208 | 10.1 | 31 | Wu and Kaiser (1995) |
| GSU1776 | *Geobacter sulfurreducens** (GsOxpG) | OxpG | S | 164 | 11.0 | 22 | Mehta et al. (2006) |

**Table S2.** **Characteristics of type IV pilins and pseudopilins involved in functions other than long-range electron transport, in order of increasing aromatic density.** A: attachment; C: competence; F: fibrial assembly; M: motility; S: secretion of proteins.  *****Predicted to be conductive based on the criteria used for this study.

| **IMG ID/ GenBank BioProject** | **Environment** | **Sample** | **Protein-coding genes** | **Putative e-pilins / T4aPs** | **Aromatics (%)** | **Mature length (aa)** |
| --- | --- | --- | --- | --- | --- | --- |
| PRJNA521166 | Lake Matano, Indonesia | Ferruginous  sediment | 1,157,873 | 2/93 | 12.4, 12.7 | 110, 113 |
| 3300031898, [3300031587](https://img.jgi.doe.gov/cgi-bin/m/main.cgi?section=TaxonDetail&page=taxonDetail&taxon_oid=3300031587), [3300031877](https://img.jgi.doe.gov/cgi-bin/m/main.cgi?section=TaxonDetail&page=taxonDetail&taxon_oid=3300031877) | Lake Towuti, Indonesia |  | 10,203,152 | 3/113 | 11.3, 14.4 | 115, 118 |
| 3300014720, 3300013122, 3300013123, 3300013125, 3300013126, 3300013131, 3300013132, 3300013136, 3300013137, 3300013138 | Kabuno Bay, Lake Kivu,  Democratic Republic of Congo | Ferruginous  water column | 24,761,074 | 50/1,118 | 9.8-13.8 | 58-162 |
| 3300027975 | Prairie Pothole Lake, North Dakota, USA | Sediment | 1,493,055 | 0/24 | N/A | N/A |
| 3300025843 | Lake Baikal, Russia |  | 1,180,048 | 0/32 | N/A | N/A |
| 3300019765 | Broad Kill River, Delaware, USA |  | 669,074 | 0/1 | N/A | N/A |
| 3300014656 | Aspo, Sweden | Groundwater | 3,495,273 | 3/140 | 10.0-11.9 | 60-146 |
| PRJNA297582, PRJNA362739 | Crystal Geyser, Utah, USA |  | 1,537,765 | 10/413 | 10.5-12.5 | 111-126 |
| PRJNA273161 | Rifle, Colorado, USA |  | 633,116 | 1/389 | 15.5 | 155 |
| PRJNA321556 | Horonobe, Japan |  | 557,865 | 6/37 | 10.9-12.3 | 64-146 |
| 3300020183, 3300021092 | Lake Tanganyika, Africa | Water column | 4,124,017 | 0/13 | N/A | N/A |
| 3300021437 | McNutts Creek, Athens, Georgia, USA |  | 464,898 | 1/5 | 10.1 | 129 |
| 3300024317 | Indiana, USA | Forest soil | 530,797 | 0/12 | N/A | N/A |
| 3300028037 | Norway |  | 161,101 | 0/0 | N/A | N/A |
| 3300023019 | Czech Republic |  | 80,325 | 0/0 | N/A | N/A |
| PRJNA391950 | North Pond, Mid-Atlantic Ridge | Crustal aquifer | 494,807 | 4/43 | 9.9-13.8 | 131-146 |
|  |  |  | **Total** | **99/2,433** | **9.8-17.2** | **58-162** |
| SAMN10492640 | Lake Matano Enrichment | 490-d Fe/CH_4_ (311FMe) | 66,924 | 1/4 | 10.9 | 64 |
| SAMN10411764 | Lake Matano Enrichment | 490-d Fe/N_2_ (311FN) | 268,143 | 5/16 | 10.9-15.3 | 58-108 |
| PRJNA489678 | Lake Matano Enrichment | 395-d Mn/N_2_ (s9) | 7,050 | 0/2 | N/A | N/A |
| PRJNA489678 | Lake Matano Enrichment | 395-d Mn/N_2_ (s11) | 45,534 | 0/3 | N/A | N/A |
|  |  |  | **Total** | **6/25** | **10.9-15.3** | **58-108** |

**Table S3. Putative e-pilin genes identified from environmental metagenomes and long-term enrichment of Lake Matano sediment in the presence of metal oxides.** Putative e-pilins are defined as containing (≥9.8% aromatic amino acids, ≤22-aa aromatic gaps, and aromatic amino acids at residues 1, 24, 27, 50 and/or 51, and 32 and/or 57. See **Supplemental Data File** for sequence and details about each putative e-pilin.

**Figure S1. Putative operon with multiple extracellular multiheme cytochromes found in *Candidatus* Omnitrophica bacterium CG11_big_fil_rev_8_21_14_0_20_42_13.** Numbers represent number of cytochromes in each gene product. Predicted protein locations based on psortb analysis. E: extracellular; OM: outer membrane; P: periplasmic; C: cytoplasmic; CM: cytoplasmic membrane. NCBI accession numbers for each protein (PIQ89515-PIQ89529) are labeled beneath each gene.

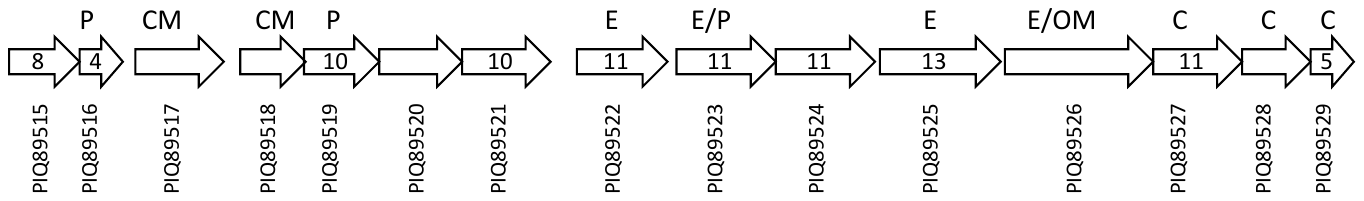

**Supplemental References**

Arrieta, J.G., Sotolongo, M., Menéndez, C., Alfonso, D., Trujillo, L.E., Soto, M. et al. (2004) A type II protein secretory pathway required for levansucrase secretion by *Gluconacetobacter diazotrophicus*. *J Bacteriol* **186**: 5031-5039.

Bakaletz, L.O., Baker, B.D., Jurcisek, J.A., Harrison, A., Novotny, L.A., Bookwalter, J.E. et al. (2005) Demonstration of type IV pilus expression and a twitching phenotype by *Haemophilus influenzae*. *Infect Immun* **73**: 1635-1643.

Bhaya, D., Bianco, N.R., Bryant, D., and Grossman, A. (2000) Type IV pilus biogenesis and motility in the cyanobacterium *Synechocystis* sp. PCC6803. *Mol Microbiol* **37**: 941-951.

Burrows, L.L. (2012) *Pseudomonas aeruginosa* twitching motility: type IV pili in action. *Annual Review of Microbiology* **66**: 493-520.

Chanyi, R.M., and Koval, S.F. (2014) Role of type IV pili in predation by *Bdellovibrio bacteriovorus*. *PloS One* **9**: e113404.

Chen, I., Provvedi, R., and Dubnau, D. (2006) A macromolecular complex formed by a pilin-like protein in competent *Bacillus subtilis*. *J Biol Chem* **281**: 21720-21727.

Doig, P., Todd, T., Sastry, P.A., Lee, K., Hodges, R.S., Paranchych, W., and Irvin, R. (1988) Role of pili in adhesion of *Pseudomonas aeruginosa* to human respiratory epithelial cells. *Infect Immun* **56**: 1641-1646.

Dougherty, B.A., and Smith, H.O. (1999) Identification of *Haemophilus influenzae* Rd transformation genes using cassette mutagenesis. *Microbiology* **145**: 401-409.

Essex-Lopresti, A.E., Boddey, J.A., Thomas, R., Smith, M.P., Hartley, M.G., Atkins, T. et al. (2005) A type IV pilin, PilA, contributes to adherence of *Burkholderia pseudomallei* and virulence *in vivo*. *Infect Immun* **73**: 1260-1264.

Graupner, S., Frey, V., Hashemi, R., Lorenz, M.G., Brandes, G., and Wackernagel, W. (2000) Type IV pilus genes pilA and pilC of *Pseudomonas stutzeri* are required for natural genetic transformation, and pilA can be replaced by corresponding genes from nontransformable species. *J Bacteriol* **182**: 2184-2190.

Hahn, H.P. (1997) The type-4 pilus is the major virulence-associated adhesin of *Pseudomonas aeruginosa*–a review. *Gene* **192**: 99-108.

Han, X., Kennan, R.M., Parker, D., Davies, J.K., and Rood, J.I. (2007) Type IV fimbrial biogenesis is required for protease secretion and natural transformation in *Dichelobacter nodosus*. *J Bacteriol* **189**: 5022-5033.

Howard, S.P., Critch, J., and Bedi, A. (1993) Isolation and analysis of eight exe genes and their involvement in extracellular protein secretion and outer membrane assembly in *Aeromonas hydrophila*. *J Bacteriol* **175**: 6695-6703.

Kang, Y., Liu, H., Genin, S., Schell, M.A., and Denny, T.P. (2002) *Ralstonia solanacearum* requires type 4 pili to adhere to multiple surfaces and for natural transformation and virulence. *Mol Microbiol* **46**: 427-437.

Kehl-Fie, T.E., Miller, S.E., and Geme, J.W.S. (2008) *Kingella kingae* expresses type IV pili that mediate adherence to respiratory epithelial and synovial cells. *J Bacteriol* **190**: 7157-7163.

Laurenceau, R., Péhau-Arnaudet, G., Baconnais, S., Gault, J., Malosse, C., Dujeancourt, A. et al. (2013) A type IV pilus mediates DNA binding during natural transformation in *Streptococcus pneumoniae*. *PLoS Pathogens* **9**: e1003473.

Li, Y., Hao, G., Galvani, C.D., Meng, Y., De La Fuente, L., Hoch, H., and Burr, T.J. (2007) Type I and type IV pili of *Xylella fastidiosa* affect twitching motility, biofilm formation and cell–cell aggregation. *Microbiology* **153**: 719-726.

Liu, X., Wang, S., Xu, A., Zhang, L., Liu, H., and Ma, L.Z. (2019) Biological synthesis of high-conductive pili in aerobic bacterium *Pseudomonas aeruginosa*. *Appl Micro Biotechnol* **103**: 1535-1544.

Marrs, C.F., Schoolnik, G., Koomey, J.M., Hardy, J., Rothbard, J., and Falkow, S. (1985) Cloning and sequencing of a *Moraxella bovis* pilin gene. *J Bacteriol* **163**: 132-139.

Mehta, T., Childers, S.E., Glaven, R., Lovley, D.R., and Mester, T. (2006) A putative multicopper protein secreted by an atypical type II secretion system involved in the reduction of insoluble electron acceptors in *Geobacter sulfurreducens*. *Microbiology* **152**: 2257-2264.

Nassif, X., Beretti, J.-L., Lowy, J., Stenberg, P., O'Gaora, P., Pfeifer, J. et al. (1994) Roles of pilin and PilC in adhesion of *Neisseria meningitidis* to human epithelial and endothelial cells. *Proc Natl Acad Sci* **91**: 3769-3773.

Nien-Tai, H., Wei-Ming, L., Meng-Shiunn, L., Avon, C., Shu-Chung, C., Yu-Ling, S., and Ling-Yun, C. (2002) XpsG, the major pseudopilin in *Xanthomonas campestris* pv. *campestris*, forms a pilus-like structure between cytoplasmic and outer membranes. *Biochem J* **365**: 205-211.

Paranjpye, R.N., and Strom, M.S. (2005) A *Vibrio vulnificus* type IV pilin contributes to biofilm formation, adherence to epithelial cells, and virulence. *Infect Immun* **73**: 1411-1422.

Pugsley, A.P. (1993) Processing and methylation of PulG, a pilin‐like component of the general secretory pathway of *Klebsiella oxytoca*. *Mol Microbiol* **9**: 295-308.

Rakotoarivonina, H., Larson, M.A., Morrison, M., Girardeau, J.-P., Gaillard-Martinie, B., Forano, E., and Mosoni, P. (2005) The *Ruminococcus albus* pilA1–pilA2 locus: expression and putative role of two adjacent pil genes in pilus formation and bacterial adhesion to cellulose. *Microbiology* **151**: 1291-1299.

Reguera, G., McCarthy, K.D., Mehta, T., Nicoll, J.S., Tuominen, M.T., and Lovley, D.R. (2005) Extracellular electron transfer via microbial nanowires. *Nature* **435**: 1098-1101.

Rossier, O., Starkenburg, S.R., and Cianciotto, N.P. (2004) *Legionella pneumophila* type II protein secretion promotes virulence in the A/J mouse model of Legionnaires' disease pneumonia. *Infect Immun* **72**: 310-321.

Rudel, T., van Putten, J.P., Gibbs, C.P., Haas, R., and Meyer, T.F. (1992) Interaction of two variable proteins (PilE and PilC) required for pilus‐mediated adherence of *Neisseria gonorrhoeae* to human epithelial cells. *Mol Microbiol* **6**: 3439-3450.

Rumszauer, J., Schwarzenlander, C., and Averhoff, B. (2006) Identification, subcellular localization and functional interactions of PilMNOWQ and PilA4 involved in transformation competency and pilus biogenesis in the thermophilic bacterium *Thermus thermophilus* HB27. *FEBS J* **273**: 3261-3272.

Sandkvist, M., Michel, L.O., Hough, L.P., Morales, V.M., Bagdasarian, M., Koomey, M., and DiRita, V.J. (1997) General secretion pathway (eps) genes required for toxin secretion and outer membrane biogenesis in *Vibrio cholerae*. *J Bacteriol* **179**: 6994-7003.

Sauvonnet, N., Vignon, G., Pugsley, A.P., and Gounon, P. (2000) Pilus formation and protein secretion by the same machinery in Escherichia coli. *EMBO J* **19**: 2221-2228.

Stone, B.J., and Kwaik, Y.A. (1998) Expression of multiple pili by *Legionella pneumophila*: identification and characterization of a type IV pilin gene and its role in adherence to mammalian and protozoan cells. *Infect Immun* **66**: 1768-1775.

Tan, Y., Adhikari, R.Y., Malvankar, N.S., Ward, J.E., Woodard, T.L., Nevin, K.P., and Lovley, D.R. (2017) Expressing the *Geobacter metallireducens* PilA in *Geobacter sulfurreducens* yields pili with exceptional conductivity. *mBio* **8**: e02203-02216.

Tan, Y., Adhikari, R.Y., Malvankar, N.S., Ward, J.E., Nevin, K.P., Woodard, T.L. et al. (2016) The low conductivity of *Geobacter uraniireducens* pili suggests a diversity of extracellular electron transfer mechanisms in the genus *Geobacter*. *Front Microbiol* **7**: 980.

Thormann, K.M., Saville, R.M., Shukla, S., Pelletier, D.A., and Spormann, A.M. (2004) Initial phases of biofilm formation in *Shewanella oneidensis* MR-1. *J Bacteriol* **186**: 8096-8104.

Tønjum, T., Marrs, C.F., Rozsa, F., and Bøvre, K. (1991) The type 4 pilin of *Moraxella nonliquefaciens* exhibits unique similarities with the pilins of *Neisseria gonorrhoeae* and *Dichelobacter* (*Bacteroides*) *nodosus*. *Microbiology* **137**: 2483-2490.

Villar, M.T., Hirschberg, R.L., and Schaefer, M.R. (2001) Role of the *Eikenella corrodens* pilA locus in pilus function and phase variation. *J Bacteriol* **183**: 55-62.

von Tils, D., Blädel, I., Schmidt, M.A., and Heusipp, G. (2012) Type II secretion in *Yersinia*—a secretion system for pathogenicity and environmental fitness. *Front Cell Infect Microbiol* **2**: 160.

Walker, D.J., Nevin, K.P., Holmes, D.E., Rotaru, A.-E., Ward, J.E., Woodard, T.L. et al. (2019a) *Syntrophus* conductive pili demonstrate that common hydrogen-donating syntrophs can have a direct electron transfer option. *BioRXiv*: 479683.

Walker, D.J.F., Martz, E., Holmes, D.E., Zhou, Z., Nonnenmann, S.S., and Lovley, D.R. (2019b) The archaellum of *Methanospirillum hungatei* is electrically conductive. *mBio* **10**: e00579-00519.

Walker, D.J.F., Adhikari, R.Y., Holmes, D.E., Ward, J.E., Woodard, T.L., Nevin, K.P., and Lovley, D.R. (2018) Electrically conductive pili from pilin genes of phylogenetically diverse microorganisms. *ISME J* **12**: 48–58.

Wu, S.S., and Kaiser, D. (1995) Genetic and functional evidence that type IV pili are required for social gliding motility in *Myxococcus xanthus*. *Mol Microbiol* **18**: 547-558.

Zhang, Y., Tennent, J.M., Ingham, A., Beddome, G., Prideaux, C., and Michalski, W.P. (2000) Identification of type 4 fimbriae in *Actinobacillus pleuropneumoniae*. *FEMS Microbiol Lett* **189**: 15-18.
